# The assembly of wild natural isolates define neuronal integrity and life history traits of co-inhabiting *C. elegans*

**DOI:** 10.1101/2022.09.26.509631

**Authors:** Sebastian Urquiza-Zurich, Victor Antonio Garcia-Angulo, Paula Burdisso, M. Fernanda Palominos, Lucia Fernandez-Hubeid, Juan Pablo Castillo, Andrea Calixto

## Abstract

Bacterivore nematodes are the most abundant animals in the biosphere, largely contributing to global biogeochemistry. The effect of environmental microbes as source of associated microbiota and natural diet on their life history traits of nematodes is likely to impact the general health of the biosphere. *Caenorhabditis elegans* is a unique model to study the behavioral and physiological outputs of different available microbial diets. Nonetheless, most studies are on monoaxenic cultures of laboratory bacteria while the effect of natural microbiota isolates has only recently started to be reported. Here, we quantified physiological, phenotypical and behavioral traits of worms feeding on two bacteria that co-isolated with wild nematodes and tested how combinations of these isolates with other bacteria affected the traits measured. These bacteria were identified as a putative novel species of *Stenotrophomonas* denominated *Stenotrophomonas* sp. Iso1 and a strain of *Bacillus pumilus* designated Iso2. The isolates induced distinctive behaviors and development patterns that changed in mixes of the two bacteria and/or the pathogen *Salmonella enterica*. Focusing on the degeneration rate of the touch circuit of *C. elegans* we show that *B. pumilus* alone is protective while the mix with *Stenotrophomonas* sp. is degenerative. The analysis of the metabolite content of each isolate and their combination identified NAD+ as potentially neuroprotective. *In vivo* supplementation shows that NAD+ restores neuroprotection to the mixes and also to individual non-protective bacteria. The results highlight the need to study the physiological effects of bacteria resembling native diets in a multicomponent scenario rather than using single isolates.

**Importance:** The behavioral decisions of animals depend on their microbiota. In nature it is unknown how this interaction affects the health of the biosphere. To study how the nematode-bacteria relationship impacts the life history traits of these animals, we isolated bacteria found in association with wild nematodes and tested their influence as single species and consortia, in the life history traits of the model *C. elegans*. We identify metabolites from wild bacteria that change these traits. The bacteria isolated were identified a *Stenotrophomonas sp* and a *B. pumilus*. We find that all traits depend on the biota composition. For example, *B. pumilus* is neuroprotective to degenerating neurons of the touch circuit of *C. elegans* needed to sense and escape from predators in the wild. The co-culture with *Stenotrophomonas sp*. eliminates the protection. We identified NAD+ as the metabolite lost in the mix, and show that NAD+ by itself is neuroprotective.

## Introduction

All eukaryotes live intimately with microbial communities. Within these ecosystems, the communication with microbes substantially impacts the life history traits of the eukaryotic hosts (1, 2). However, how the native microbiota affects the life-history traits of the host and promotes adaptation is still largely unknown. The bacterivore nematode *Caenorhabditis elegans* offers a simple framework to manipulate the composition and metabolite content of its microbial diet to dissect specific molecules that direct behavior and phenotypical outputs in the animal.

*C. elegans* has been typically grown on monoaxenic cultures of the *Escherichia coli* strain OP50 (3). Feeding worms on other bacteria allowed the discovery that developmental rate depends on specific bacterial metabolites such as vitamin B12 (4) or that neurodegeneration can be delayed by gamma-aminobutyric acid (GABA) produced by *E. coli* HT115 (5). Those pioneer works however were also done with monocultures of laboratory bacteria opening the question of how does complex natural microbiota influence behavior and physiology. We hypothesize that co-existing consortia of environmental bacteria promote behavioral signatures different from each individual bacterium in host *C. elegans*. The study of environmental bacteria might also serve as a platform to dissect benefits or detriments that microbial associations can cause to hosts in natural environments.

The microbiota of *C. elegans* and other Rhabditida species from Europe and North America is composed mainly by species of the *Enterobacteriaceae* family (6), *Pseudomonas, Stenotrophomonas, Xanthomonas, Ochrobactrum* and *Sphingomonas* genera (7, 8). However, nematodes from South America and their associated bacteria have not been studied. We isolated bacteria carried by wild nematodes from the soil in a semi-arid location in central Chile, identifying a potentially new species of *Stenotrophomonas* and a novel strain of *Bacillus pumilus*. Monocultures produced life-history trait output different from co-cultures. Specifically, *B. pumilus* is neuroprotective to *C. elegans* with degenerating touch circuitry while its co-culture with Stenotrophomonas eliminates the trait. Metabolomic analysis of single bacteria compared to their mix coupled to *in vivo* validation identified bacterial neuroprotective metabolites.

## Results

### Isolation and identification of microbes from wild Chilean nematodes

Adaptation to environmental change requires the work of co-inhabiting organisms, especially those most ubiquitous such as bacteria and nematodes. To reveal how bacteria in natural environments modulate life history traits of the nematode *C. elegans*, we collected nematodes from a Chilean scrubland, characterized by mountain ranges and Andean steppes with increasingly prominent hydric stress. We placed soil samples on growth plates seeded with *E. coli* OP50 lawns, the standard laboratory diet of *C. elegans*. Wild worms present in the soil sample migrated towards *E. coli* OP50 carrying wild bacteria that grew on the plates. Wild nematodes preferred to feed on these colonies for generations rather than on *E. coli* OP50. We isolated two distinctive colonies from the wild bacterial growth, representing a putative preferred native-worm diet. These isolates were denominated Iso1 and Iso2.

We sequenced the genomes of Iso 1 and Iso 2 followed by genome-based identification analysis. We employed the Type Strain Genome Server (TYGS) pipeline for the identification of species (**Fig. 1**). This was complemented by the determination of the average nucleotide identity (ANI) between the isolate genomes and their closest type strain detected by TYGS, using OrthoANIu (9). For Iso1, the closest type was *Stenotrophomonas humi* DSM 18929. Nonetheless, both the 16S-based and the GBDP-based phylogram (**Fig. 1A** and **B**) locate the isolate in a different species and subspecies cluster to this type strain. The dDDH analysis renders a d_4_ value of 37.8% of digital hybridization with the genome of *S. humi* DSM 18929. In the dDDH procedure, the d_4_ value under 70% indicates separated species (10) (**Table 1**). The 37.8% dDDH with a confidence interval of 35.4 -40.3 for the comparison against *S. humi* DSM 18929 places the isolate well beyond the threshold to be considered a new species. Moreover, the OrhoANIu analysis determined an ANI of 89.5% for the comparison of Iso1 against *S. humi* DSM 18929 (ANI value over 95-97% indicates strains belonging to the same species). Thus, even using the loose 95% ANI threshold, this result supports that Iso1 is a novel species within the *Stenotrophomonas* genus. Hence, this isolate was denominated *Stenotrophomonas* sp. Iso1.

**Figure 1.**
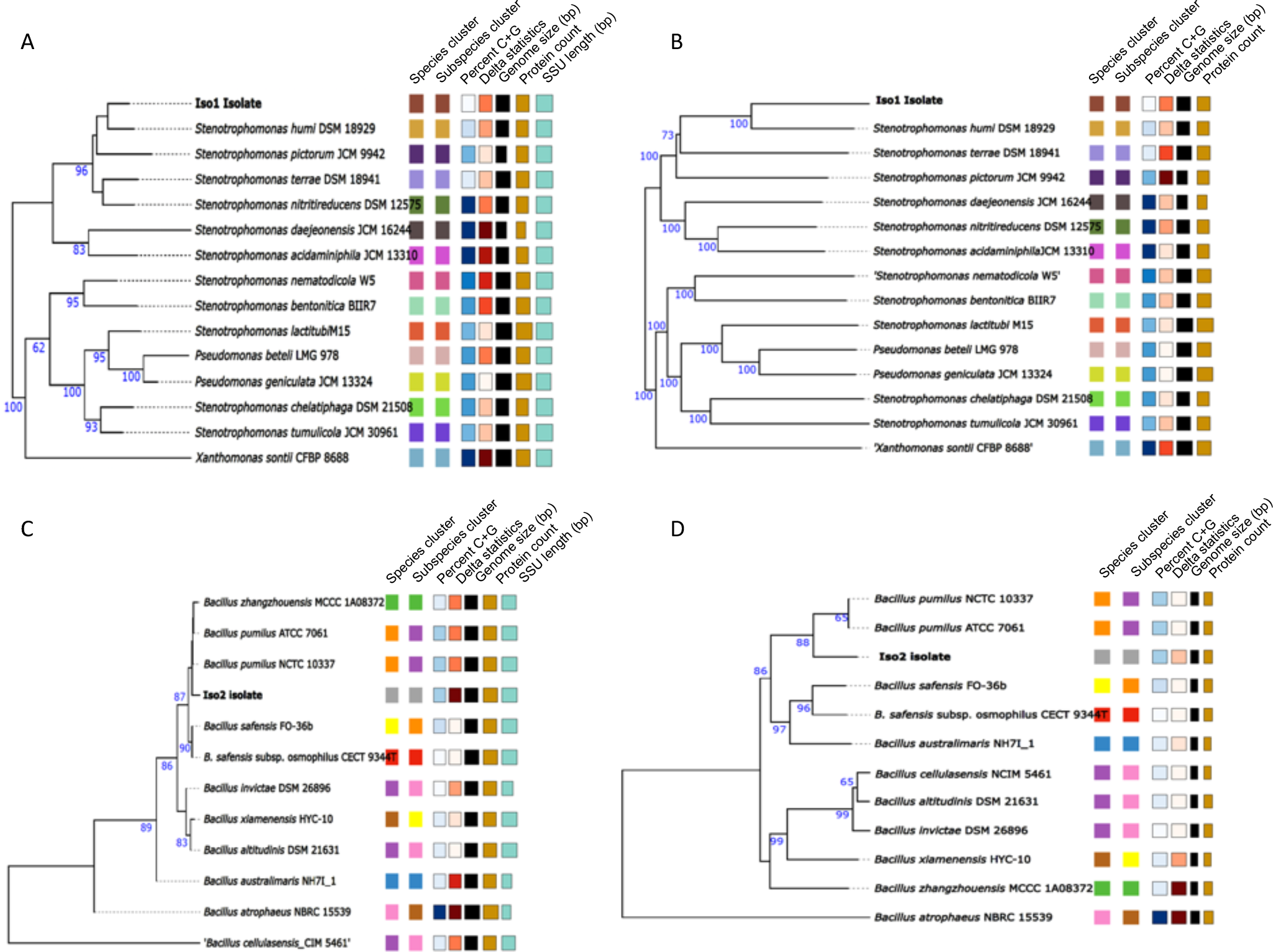
TYGS phylograms results for the bacterial isolates. 16S-based phylogram (**A** and **C**) and GBDP-based phylogram (**B** and **D**) for Iso1 and Iso2, respectively. In both cases the phylograms analysis includes the closest type strains in the databases. The color of the squares group strains belonging to the same cluster as defined by each of the parameters indicated at the top of the squares. SSU: small subunit rRNA.

**Table 1.**
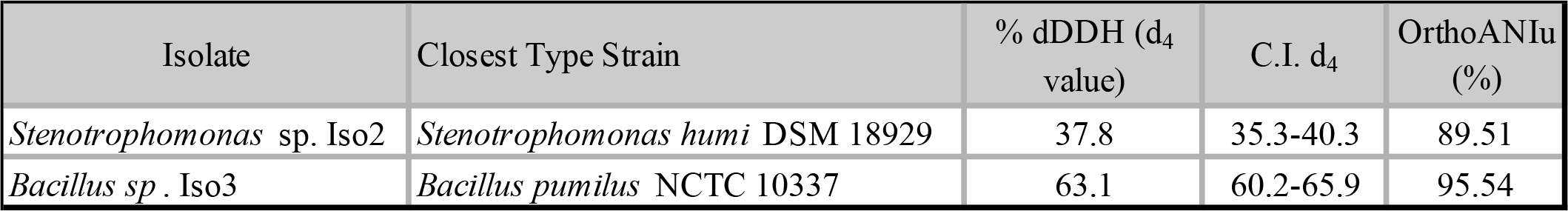

The 16S-based and GBDP-based phylograms **(Fig. 1C** and **D**) place Iso2 in a different species and subspecies cluster to that of its closest type strain, *Bacillus pumilus* NCTC 10337. Nonetheless, the obtained dDDH value against this type strain is 63.1% with a 60.2-65.9% C.I (**Table 1)**. These values are close to the 70% threshold. Moreover, the OrhoANIu value of Iso2 against this species is 95.54, which would indicate that the isolate is a strain of *B. pumilus*. Thus, it is not clear that this bacterium represents a novel species separated from *B. pumilus* hence we it as designated *Bacillus pumilus* Iso2.

### *C. elegans* dietary choice of natural isolates and their consortia is bacterial context dependent

*Stenotrophomonas* sp. Iso1 and *B. pumilus* Iso2 are likely part of the natural diet of wild nematodes. We tested the preference of *C. elegans* for the isolates, individually or in consortia. The alternative choices were *E. coli* OP50, or *E. coli* HT115. The test also included *S. enterica*, which can infect and cause disease in worms, thus representing a detrimental diet (11). While *Stenotrophomonas sp*. was strongly preferred over *E. coli* OP50 at both time points, *B. pumilus* induced a less marked preference that was evident only at 12 hours (**Fig. 2A**). In contrast, *S. enterica* caused worms to strongly prefer *E. coli* OP50 (**Fig. 2A**). These patterns changed when bacteria were mixed. The mix of the two wild isolates causes nematodes to prefer *E. coli* OP50, while the mixes of either wild isolate with *S. enterica* were preferred, being the mix with *B. pumilus* the most selected (**Fig. 2A)**. Finally, the mix of the three bacteria, caused *C. elegans* to prefer *E. coli* OP50. This suggests that animals might prefer initially what is familiar to them (12). A different scenario is observed when *E. coli* HT115-new to the animals-, is used as alternative. Animals strongly prefer *E. coli* HT115 over *S. enterica* or *B. pumilus*, while *Stenotrophomonas* sp. was chosen equally to *E. coli* HT115. However, when *Stenotrophomonas* sp. was mixed with *S. enterica* or *B. pumilus*, animals preferred *E. coli* HT115; and the mix of *S. enterica* with *B. pumilus* was selected equally as *E. coli* HT115. Notably, contrary to the experiment with *E. coli* OP50, the mix of the three bacteria was markedly preferred over *E. coli* HT115 (**Fig. 2B**). These indicate that food choice is context dependent and that in addition to novelty vs familiarity, worms might sense molecules present only in the co-cultures that direct their preference.

**Figure 2.**
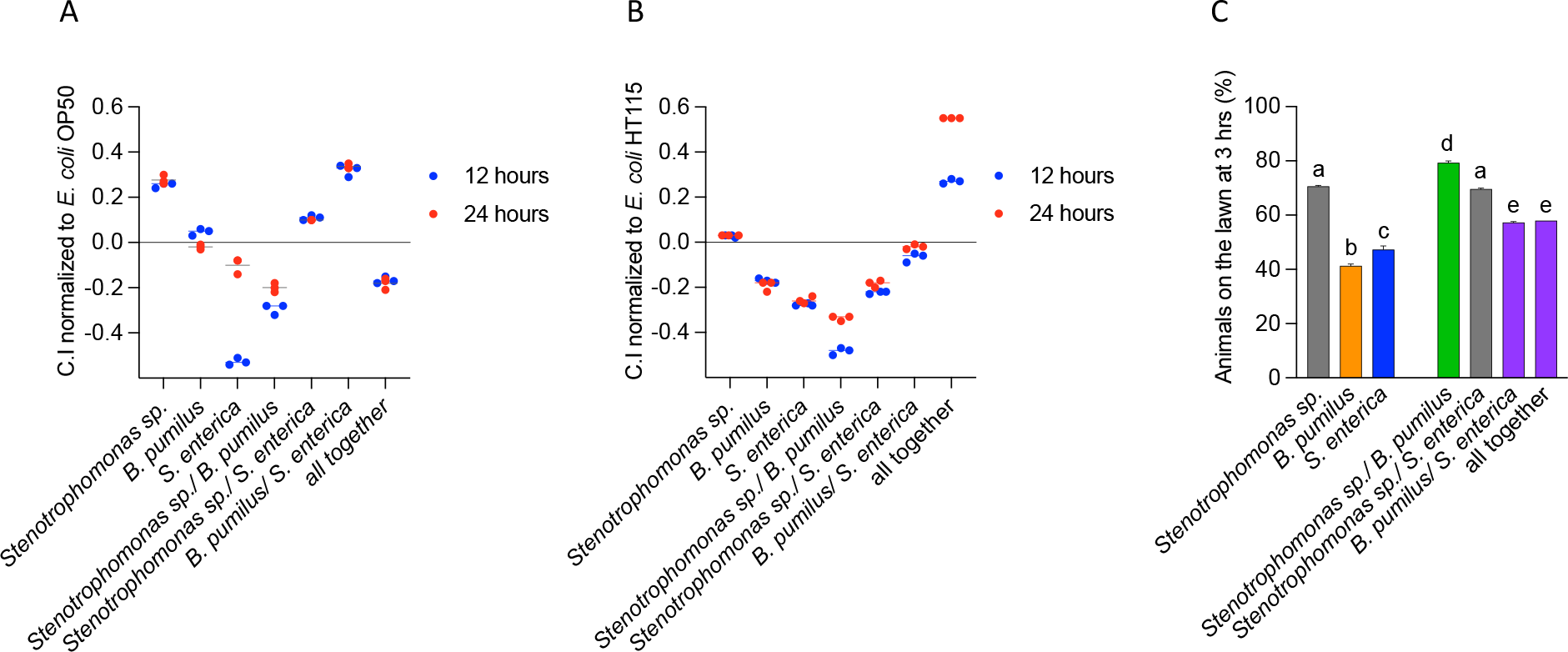
*C. elegans* preference of bacteria and consortia. **A-B**. Chemotaxis index (formula) of each individual bacteria or mixes normalized to E. coli OP50 (**A**) or E. coli HT115 (B) at 12 and 24 hours after exposure to food choice. **C**. Percentage of animals inside individual bacterial lawn or mixes after 3 hours of placing them in the center of the lawn.

We next examined the tendency of animals to remain in lawns, or to migrate outside after one, two and three hours. The lawn with the highest occupancy was *Stenotrophomonas* sp. with 70% at 3 hours (**Fig. 2C**), while *S. enterica* and *B. pumilus* retained under 50% occupancy (the complete quantification is on **Fig. S1**). The mix of the two wild isolates retained over 80% of animals indicating that when not competing to *E. coli* OP50, their mix is highly liked. The mix of *S. enterica* with *B. pumilus* or with the two wild isolates resulted in higher occupancies than those of *S. enterica* or *B. pumilus* alone. In this experiment, mixing *S. enterica* with *Stenotrophomonas* sp. did not affect the high worm retention produced by *Stenotrophomonas sp*.

We next asked whether nematodes dwell or roam on the bacterial lawns. Animals dwelled on *E. coli* HT115 and roamed in *E. coli* OP50, as reported before for good and mediocre diets respectively (13). Animals dwelled on *B. pumilus* but roamed on *Stenotrophomonas sp*., and *S. enterica* (**Table 2**). The mix of *B. pumilus* and *Stenotrophomonas sp*. caused dwelling. The mix of *S. enterica* with *B. pumilus* or *B. pumilus-Stenotrophomonas* sp. switches animals to roaming, suggesting that the locomotion pattern of animals is sensitive to changes in bacterial composition.

**Table 2.**

| Behavior in the bacterial lawn |  |  |
| --- | --- | --- |
|  | Dwelling | Roaming |
| *E. coli* OP50 |  | ● |
| *E. coli* HT115 | ● |  |
| *S. enterica* |  | ● |
| *Stenotrophomonas* sp. |  | ● |
| *B. pumilus* | ● |  |
| *S. enterica*/ *Stenotrophomonas* sp. |  | ● |
| *S. enterica*/ *B. pumilus* |  | ● |
| *Stenotrophomonas* sp./ *B. pumilus* | ● |  |
| all together |  | ● |

### B. pumilus accelerates pharyngeal pumping and defecation rate

As a measure of feeding rate, we measured the number of contractions of the pharynx per minute (14), and the defecation rate, a rhythm coupled to peristaltic movements caused by feeding. *B. pumilus* generates the highest number of pumps per minute (ppm) with a mean of 327, followed by *E. coli* HT115 (mean 293). *Stenotrophomonas* sp. slowed the pharyngeal pumping to 223 ppm, the lowest rate observed (**Fig. 3A**). The ppm produced by the mix of wild isolates resembled the average between the two individual bacteria, as did the mix with *S. enterica*, suggesting that each bacterium influences the pumping rate equally.

**Figure 3.**
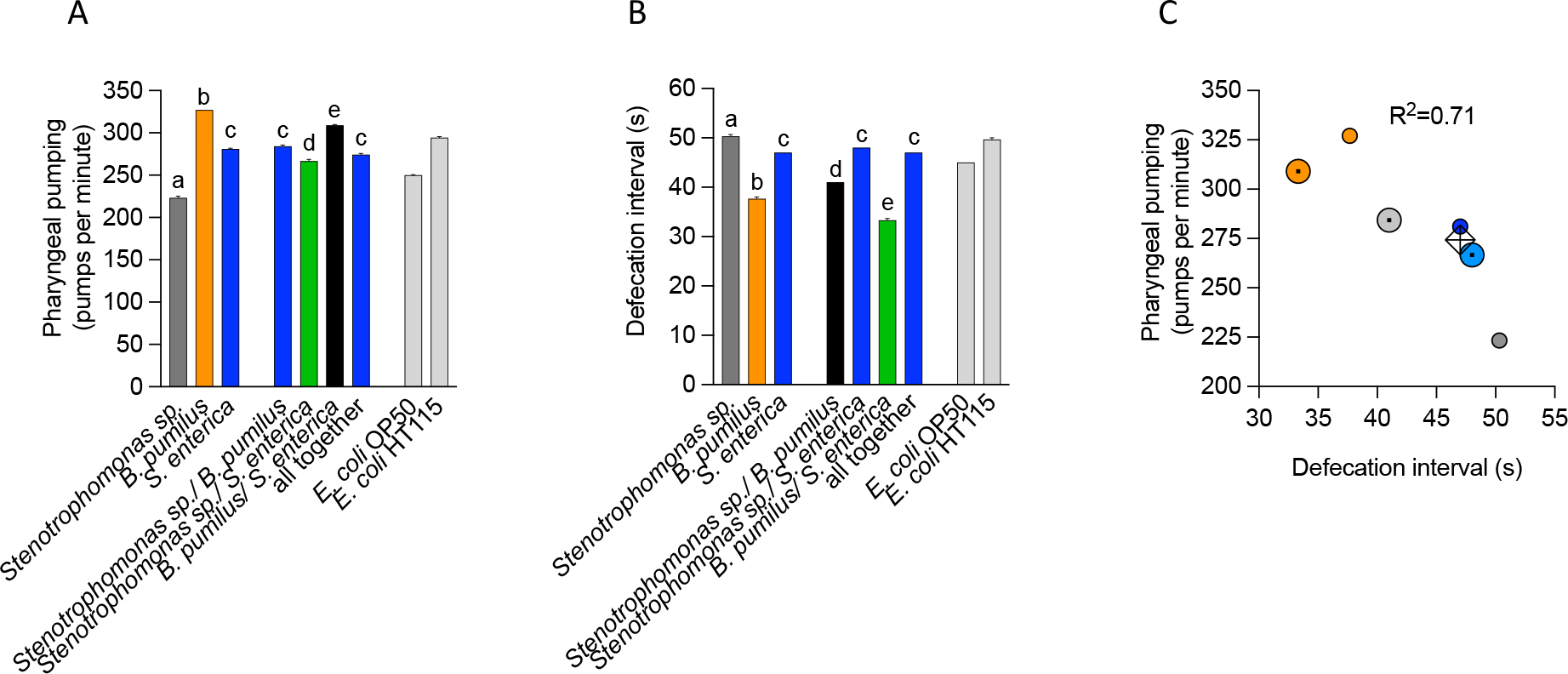
Pharyngeal pumping and defecation rates in different bacteria and consortia. **A.** Pharyngeal pumping rate (pumps per minute) of L4 animals feeding on single bacteria or consortia for one generation. *E. coli* OP50 and *E. coli* HT115 serve as control reference and were performed along the other quantifications. **B**. Defecation rate (anal contractions per minute) of L4 animals feeding on single bacteria or consortia for one generation. *E. coli* OP50 and *E. coli* HT115 serve as control reference and were performed along the other quantifications. **C**. Correlation between pharyngeal pumping rate and defecation rate using the Pearson test. Small circles are individual bacterium (orange *B. pumilus*; gray *Stenotrophomonas sp*.; blue *S. enterica*), larger circles with a dot are consortia of two bacteria (orange *S. enterica*/ *B. pumilus*; gray *Stenotrophomonas sp*./ *B. pumilus*; blue *Stenotrophomonas sp*/ *S. enterica)*, the diamond represents the mix of all bacteria.

*C. elegans* defecate every 45 seconds on *E. coli* OP50 (ref. 15, **Fig. 3B**). *B. pumilus* alone accelerates the defecation rate to a mean of 37 seconds, which is also coherent with the increased pumping rate. *Stenotrophomonas sp*. (mean 50) and *S. enterica* (mean 47) were similar to the *E. coli* controls (**Fig. 3B**). *B. pumilus* mixed with *S. enterica* generated a shorter defecation interval than *B. pumilus* alone (mean 33) and *Stenotrophomonas* sp. with *B. pumilus* (mean 41) averaged the individual isolates (**Fig. 3B**). We correlated the rate of defecation and pharyngeal pumping using a Pearson test (**Fig. 3C**) and found an inverse correlation where accelerated pumping implies a shorter defecation time.

### Wild isolates delay development into adults

We measured the developmental rate of *C. elegans* feeding on the isolates. Animals were fed with individual and co-cultures of bacteria, and the number of nematodes in each larval and adult stage was quantified every 24 hours for three days. In the first 48 hours, most animals developed similarly regardless of the diet (**Fig. S2**). At 72 hours, a delay in growth is observed on *B. pumilus* alone and when mixed with either *Stenotrophomonas sp*. or *S. enterica* compared with other mixes also containing these two bacteria (**Fig. 4A** and **B**). Because pharyngeal pumping is high on *B. pumilus* (**Fig. 3A**), it is unlikely this is due to low ingestion. An alternative is that that *B. pumilus* lacks metabolites that accelerate development such as vitamin B12 (16). *C. elegans* acyl-CoA dehydrogenase (*acdh-1)* transcripts are low in presence of vitamin B12 (4) and high in *E. coli* OP50 which does not produce the vitamin. *acdh-1::gfp* expressing animals fed with the isolates caused GFP fluorescence to be high in the intestine (**Fig. 4C** and **D**). This shows that both wild isolates have high levels of *acdh-1*, likely suggesting they lack vitamin B12.

**Figure 4.**
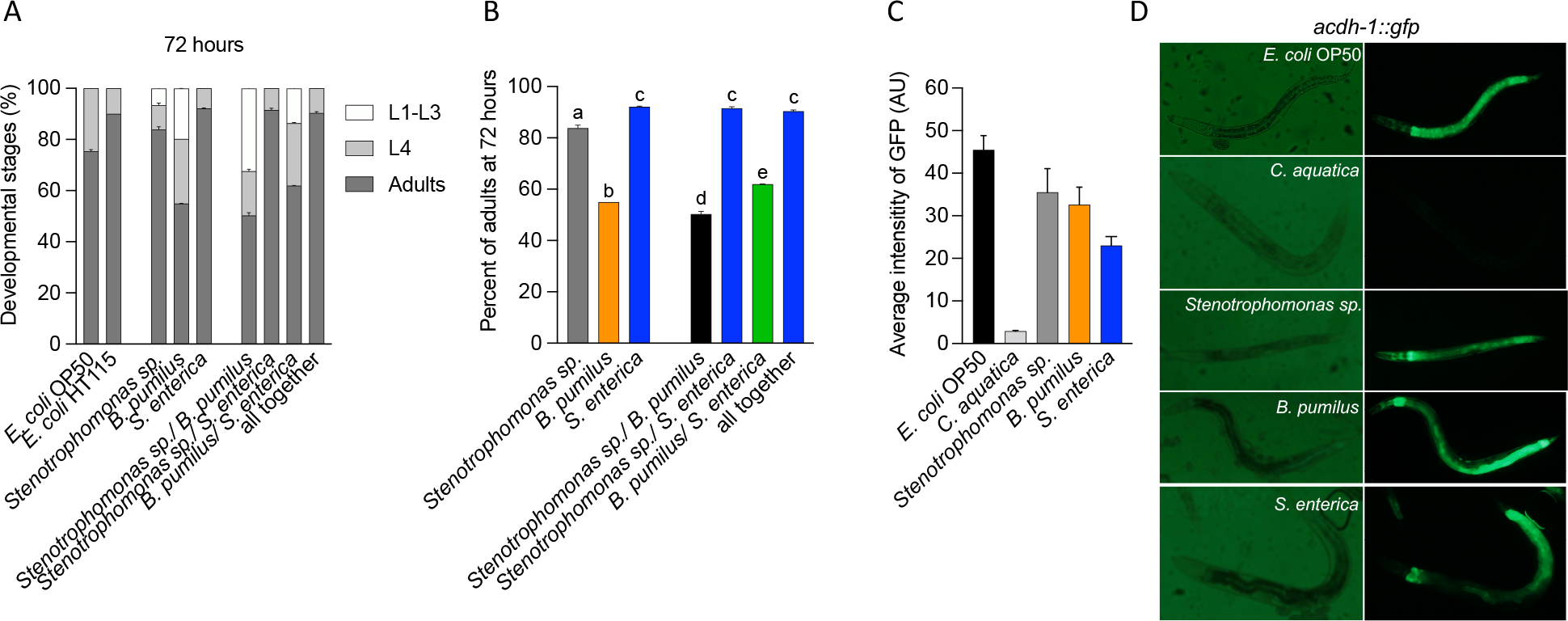
Development of animals in different bacteria and consortia. **A-B**. Percentage of animals in larval or adult stages (**A**) or only adults (**B**) at 72 hours after exposure to single bacteria or consortia. **C**. Average intensity of *acdh-1:: gfp* transgene expressed in intestine of *C. elegans* feeding on single bacteria. **D**. Representative images of **C**.

### *B. pumilus* neuroprotective effect is lost when mixed with *Stenotrophomonas* sp

In *C. elegans*, GABA producing bacteria prevent the genetically-induced neurodegeneration of the touch receptor neurons (TRN) (5). The TRNs are responsible for sensing gentle touch in the animal (17), a response that depends on the function of MEC-4(18). A mutation that affects the extracellular domain of this protein (Val713), renders a channel defective in gating, promoting the unregulated influx of sodium leading to the degeneration of the cell (19). In these worms, termed *mec-4d*, we tested whether the wild bacterial isolates and their consortia offer protection against constitutive neurodegeneration of the anterior ventral microtubule (AVM) touch neuron (20). *B. pumilus* induce neuroprotection compared to the *E. coli* OP50 diet, while *Stenotrophomonas* sp. and *S. enterica* do not provide neuroprotection (**Fig. 5A**). AVM neuronal integrity conferred by the different bacterial diets correlated with the response to the gentle touch stimulus (**Fig. 5B**). The mix of the two wild isolates eliminates the protection provided by *B. pumilus* while the mix of *B. pumilus* with *S. enterica* does not (**Fig. 5**). These results suggest that the *B. pumilus* strain may produce metabolites with neuroprotective potential, and that the expression and/or stability of such metabolites is affected by its co-culture with *Stenotrophomonas* sp.

**Figure 5.**
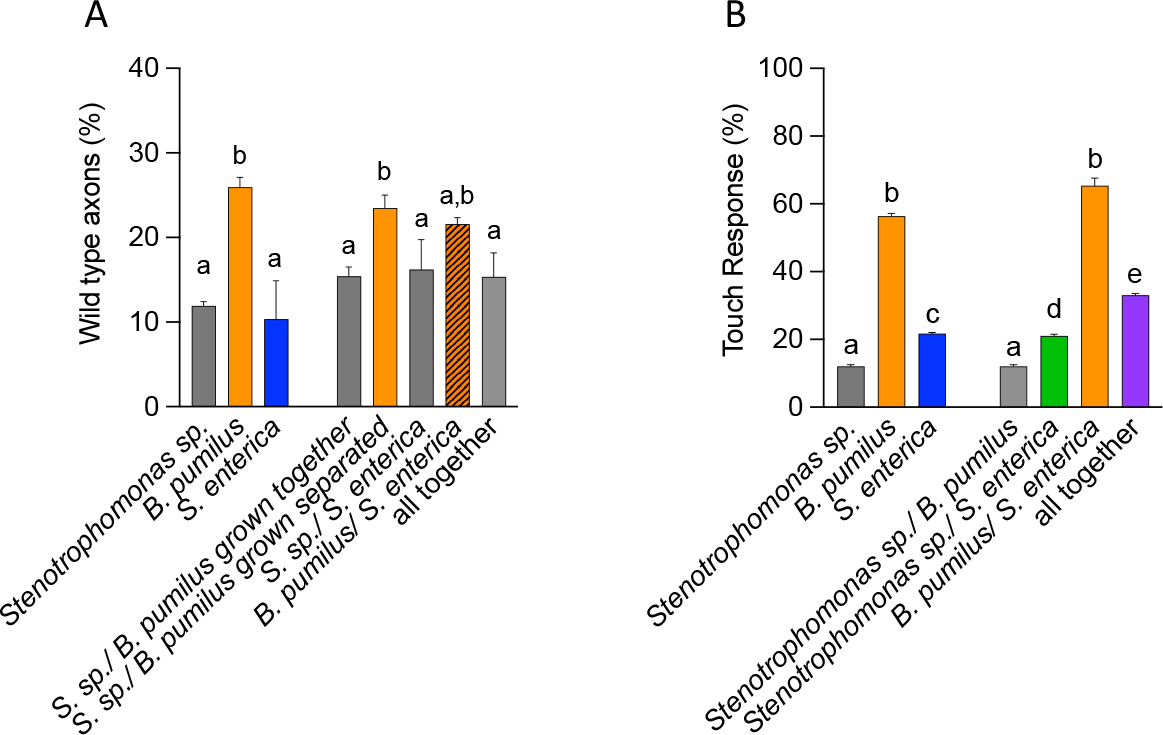
Neurodegeneration of the touch receptor neurons and response to touch in different bacteria and consortia. A-B. Percentage of wild type (protected) AVM axons (**A**) and touch response (**B**) of animals feeding on single bacteria or consortia for one generation.

### Metabolite profiling reveals NAD+ as a neuroprotector produced by native dietary bacteria

To pinpoint which bacterial metabolites underlie the neuronal protection provided by *B. pumilus*, we extracted and analyzed the global metabolite content of the isolates and their mixes by nuclear magnetic resonance (NMR) and multivariate data analysis.

We first identified patterns and trends of clustering between samples by principal component analysis (PCA) of binned ^1^H NMR datasets. In the PCA score plot (**Fig. 6A**), all samples fell within the Hotelling’s T2 ellipse at 95% confidence intervals. Three Principal Components (PC) explain 77.3% of the data variation. The main order of variation was found in PC1, which explains 45.5% of the total variation and was associated with metabolic differences between *S. enterica* relative to *B. pumilus* and *S. sp*. PC2 (22,1%), was associated with a pattern of variation between *S. sp*. and *B. pumilus*, while PC3 (9.7%) denotes different metabolic profiles between the *S. enterica* and the mixture of *S. enterica* with *B. pumilus* (**Fig. 6B-C**). Overall, the PCA revealed that the five groups of samples display different metabolic profiles, being *S. enterica* and the mixture of *S. enterica* with *S. sp*. the most distinct.

**Figure 6.**
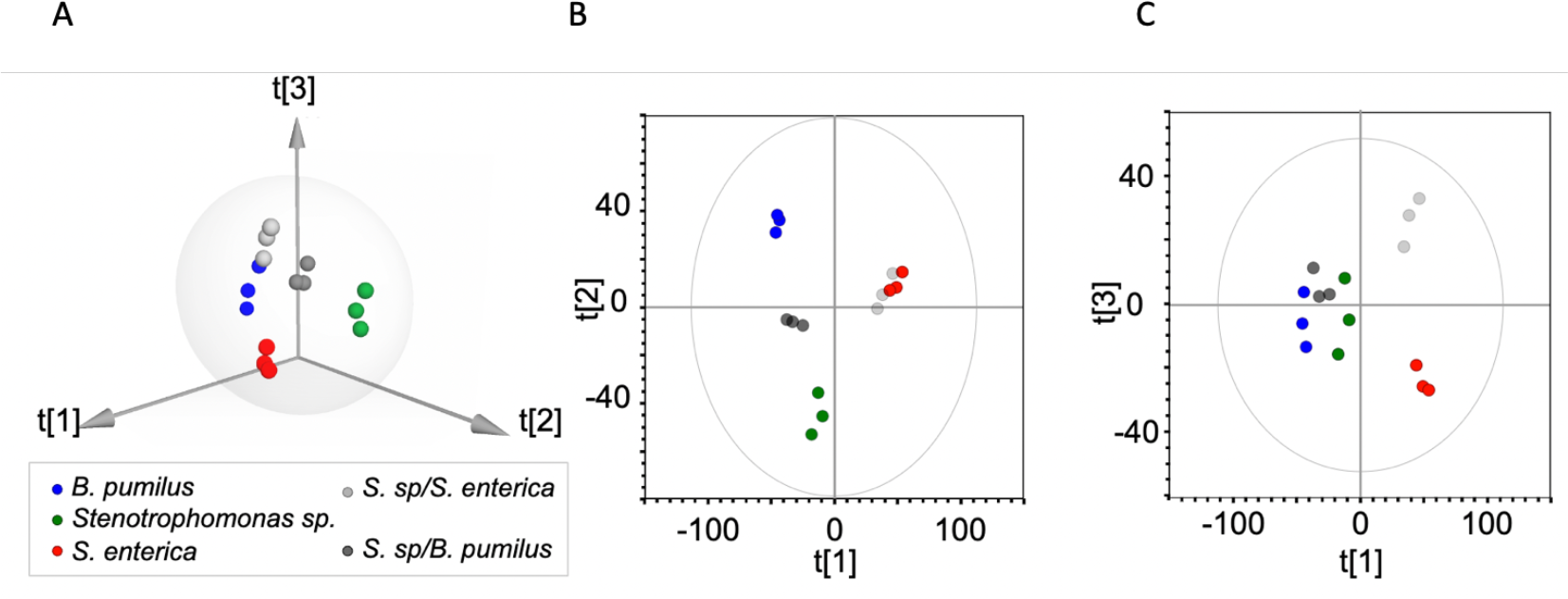
NMR Metabolomics of bacterial extracts: Exploratory data analysis by Principal Component Analysis (PCA). (A-C) Principal Component (PC) scores plot derived from 1H NMR spectra indicating clustering of bacterial strains and its mixtures. (A) 3D score plot (B and C) 2D score plot showing the first and second component (B), and the first and third one (C). Single bacterial extracts: *B. pumilus* (blue); *Stenotrophomonas sp*. (green); *S. enterica* (red); *B. pumilus/Stenotrophomonas sp*. (gray) and *B. pumilus/S. enterica* (light gray).

Neuroprotection is lost when *Stenotrophomonas sp*. and *B. pumilus* are mixed (**Fig. 5**). To identify metabolites involved in neuroprotection, we performed multivariate data analysis using the data of *S*. sp., *B. pumilus*, and the mixture of *B. pumilus* with *S. sp*. PCA was used to explore the variation within each extract. Two PCs explained 78.1% of the total variation. The split samples from the *S. sp*. and *B. pumilus* were found in the PC1 (51.1%) direction with the mixture of both located in the middle. Metabolic variation due to bacterial co-culture could be explained by PC2 (27%, **Fig. S3**).

The orthogonal projection to latent structures-discriminant analysis (OPLS-DA) method (21) was used to search for discriminant neuroprotective metabolites in *B. pumilus*. The neuroprotective class was *B. pumilus* (NP) and the neurodegenerative groups (ND) were *S. sp. And S. sp*. + *B. pumilus*. The OPLS-DA score plot (**Fig. 7A**) shows intergroup metabolic differences. The discriminant supervised model was validated by 200 permutations (**Fig. S4**). A heatmap was constructed to visualize low and highly abundant metabolites for each bacterial extract and different groups (**Fig. 7B**). Ribose and aspartate are present only in the mixed culture, while formate, alanine, nicotine amide dinucleotide (NAD+), and ethanol are present only in *B. pumilus* (**Fig. 7B**). Analysis of the Variable Importance in Projection (VIP) plot reveals differences in metabolite levels between neuroprotective and non-protective bacteria (**Fig. S5**). The metabolites with the highest VIP scores were NAD+, uracil, glycine, and ethanol.

**Figure 7.**
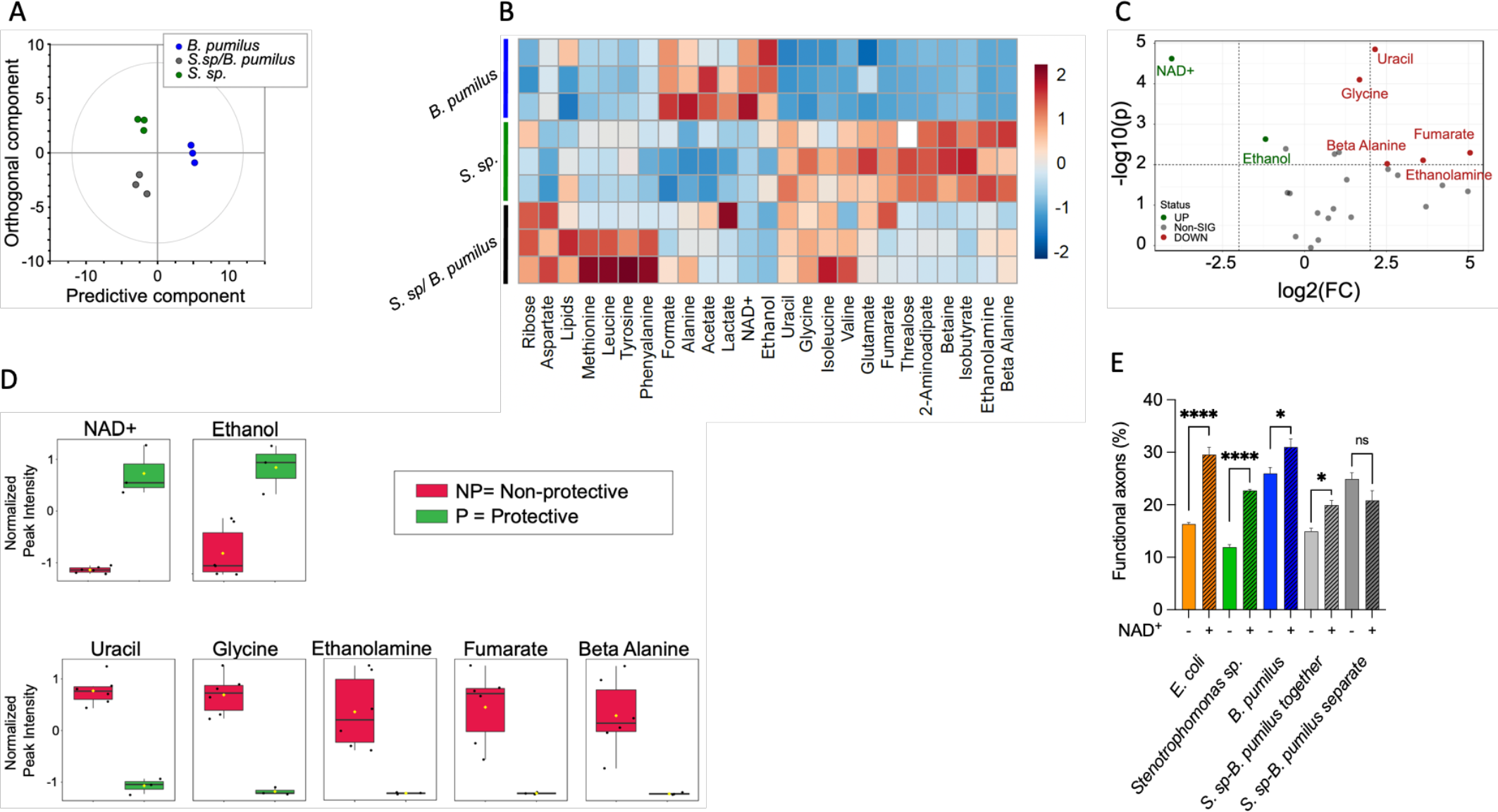
Metabolomics analysis of neuroprotective and nonprotective bacteria. Peak Intensities data matrix was normalized by PQN, mean centered and auto scaled (A) OPLS-DA score plot of protective *B. pumilus* (blue) and nonprotective *Stenotrophomonas sp*. (green) and the mixture *B. pumilus/Stenotrophomonas sp*. (gray). (B) Heatmap showing differences in relative concentrations among bacterial groups. The columns indicate different metabolites, whereas the rows indicate different samples. (C-D) Univariate metabolomics analysis. (C) Volcano plot analysis of metabolic changes in protective and nonprotective bacteria. Each point on the volcano plot was based on both p-value and fold-change values. The points which satisfy the condition p < 0.05 and fold change > 2.0 appear in red (higher levels in protective bacteria) and in blue (nonprotective bacteria). Nonsignificant markers appear in gray. (D) Box plots showed the normalized relative concentration of significant metabolites selected in volcano plot.

To focus on the independent changes in metabolite levels, univariate statistical analysis was employed to detect with higher stringency significant differences among the metabolome profiles of the groups. A Volcano plot was used to select large magnitude changes (fold change >2.0) that are also statistically significant (p<0.01) (**Fig. 7C**). This one-factor statistical analysis highlighted 7 differential metabolites (5 down and 2 up) in the NP vs. ND groups. Univariate analysis revealed ethanolamine, beta-alanine, glycine, uracil, and formic acid as potential prodegenerative metabolites whereas NAD+ and ethanol could be neuroprotective (**Fig. 7C-D**).

NAD+ was selected for *in vivo* testing (**Fig. S5**; **Fig. 7C**) on different assortments of bacteria fed to *mec-4d* animals. We observed neuroprotection in worms grown on *E. coli* OP50, *S*. sp., and *B. pumilus* supplemented with NAD+ (**Fig. 7E**), suggesting that NAD+ neuroprotection is independent of the diet and of other metabolites being synthesized. Addition of NAD+ to the co-culture of *Stenotrophomonas* sp. and *B. pumilus*, also increases the protection of *mec-4d* TRNs. When both bacteria were grown separately and then fed to the worms, the neuroprotection of *B. pumilus* is not affected, and the addition of NAD+ does not increase neuroprotection (**Fig. 7E**).

## Discussion

Microbivore nematodes are a fundamental component of the biosphere (22), and their fitness as a species is used as a parameter to test soil and niche health (23). The microbiota and microbial relationships animals establish is likely to impact the behavior of soil nematodes and therefore tightly linked to the health of these niches. Here we quantify how life history traits change as consequences of feeding animals on the natural isolates *Stenotrophomonas sp*. and *B. pumilus*, and further test how feeding on cultures with more than one bacterium affect those measurements.

We focus on more detail on the neurodegeneration of the touch circuit in *C. elegans* to study the metabolite changes that result from co-culture and that change the neuroprotective phenotype. *B. pumilus* alone is neuroprotective, but this trait is eliminated when *S. sp* and *B. pumilus* are fed together. We identify NAD+ as potentially neuroprotective and show *in vivo* that the supplementation of dietary bacteria with the chemical is strongly protective. The mix of the two isolates is only neurodegenerative when grown together in the same culture media and not when fed together to the nematodes, from independent cultures. This indicates that the dynamics of co-culture strongly affect life history traits in animals.

### Effect on life history traits of isolates and their consortia

*Stenotrophomonas* and *Bacillus* isolated with wild South American bacterivore nematodes, have been also found as part of the microbiota of nematodes elsewhere (7, 6, 24), thus likely making the association of these bacteria to the wild worms not incidental. However, whether these isolates comprise a preferred diet in the context of a natural environment, needs further assessment. Worms may be able to sense and prefer environmental bacteria that are advantageous. For example, a natural isolate of *Pantoea* that accelerates development and protects against pathogen colonization is better fitted to colonize the worm and is consistently selected as diet over other environmental strains (25).

Wild animals strongly preferred the isolates over *E. coli* OP50. *C. elegans* slightly preferred its regular *E. coli* diet at least for the first generation, which is coherent with animals preferring what is familiar to them (12). However, when offered alone, *C. elegans* remains in lawns with the isolates suggesting that in absence of familiar smells and tastes, it senses natural bacteria as a beneficial diet. The molecular mediators of this behavioral response were not explored but it is likely that metabolites produced by the bacteria are responsible for the attraction or repulsion of the nematodes. For example, *C. elegans* may regulate its food intake level depending on the nutritional quality of the diet by sensing its content of the B2 vitamin (26). Tyramine produced by an environmental *Providencia* isolate directly affects olfactory choices in the worm making it prone to keep selecting *Providencia* as food (27).

The acceleration of reproductive age together with progeny number is considered a strong indicator of fitness (28). Interestingly, *C. elegans* growing in a mix of the two isolates delays development when compared to any other diet, thus suggesting that in nature, as animals feed on a mixture of different bacterial species, the growth dynamics they experience may be slower, perhaps contributing to longevity in the wild (29).

Food availability directly modulates the rate of pharyngeal pumping (30). However, pharyngeal pumping is similar in bacteria that worms can or cannot eat (13, 31), suggesting that the pumping rate alone is not sufficient to measure food intake. We find that in both monoaxenic cultures and in mixes of *B. pumilus* with other bacteria, pharyngeal pumping rate increased, even when mixed with *Stenotrophomonas sp*., which by itself causes the slowest pumping rate. This may also imply that bacteria that are harder to ingest can induce an increase in pumping to compensate for the low food load in the digestive tract. Furthermore, we cannot rule out whether *B. pumilus* produce a molecule or metabolite that generates an increase in pharyngeal pumping (32, 33).

### Bacterial components and the neurodegeneration of the touch circuit

In the wild, nematodes use the touch circuit to escape predators such as nematophagus fungi (34). *B. pumilus* protects the neurons of *C. elegans* from constitutive genetically-induced neurodegeneration by producing the metabolite NAD+, involved in preventing oxidative stress and DNA damage in vertebrates (35, 36). We previously showed that NMAT-2 overexpression, the NAD+ producing enzyme, protects the TRNs from *mec-4d*-induced neurodegeneration (20). In this work we show that NAD+ produces by wild bacteria is relevant for neuroprotection of the host and that this protection is lost when *B. pumilus* is co-cultured with *S. sp*. While we previously found metabolites from neuroprotective laboratory *E. coli* (5), this work highlights the need to extend the search for novel neuroprotective molecules in bacteria outside the laboratory settings.

Animals fed with a combination of two or three bacteria, independently of their origin or nutritional value, change their phenotypical outcomes dramatically in comparison with when fed with individually-grown bacteria. This highlights that the understanding of the microbiota effect on animal and human physiology requires the study of complex bacterial associations.

## Materials and Methods

### Isolation of native worms and coexisting bacteria

Samples were obtained from soils pre-supplemented with fruits that rot *in situ*. Fresh, washed apples were cut and buried in the soil under a native *Schinus polygama* tree surrounded by native *Acacia caven* at the 33°22’38.3”S 70°36’54.0”W. Two weeks after, a tea spoon of soil was collected along with the decomposed apple remains. The sample was divided and deposited in three 60 mm Petri dishes containing NGM with small lawns of *E. coli* OP50. Plates were checked every 12 hours and once the nematodes appeared, they were transferred to new NGM plates with fresh *E. coli* lawns. The plates with the wild nematodes grew bacterial colonies on its *E. coli*-free surface, allowing their isolation. Bacterial colonies that contained worms feeding on them were isolates in LB and NGM media plates at 25°C.

### *C. elegans* maintenance and growth

Wild type N2 *C. elegans, mec-4d* mutants and the transgenic strain VL749 [*acdh-1p*::GFP + unc-119(+)] were grown at 20°C as previously described (3). All strains were maintained on *E. coli* OP50 prior to feeding with other bacteria. All life history traits were quantified starting with synchronized animals.

### Nematode synchronization

Larvae and adults were removed with M9 from plates full of laid embryos. Embryos remained attached to the plates and were allowed to hatch for 2 hours. 0-2 hour post hatching L1 larvae were picked with a mouth pipette in M9 and placed on experimental NGM plates seeded with the desired bacteria.

### Bacterial growth

The following strains were used as *C. elegans* diets: *E. coli* OP50, *E. coli* HT115, *S. enterica* serovar Typhimurium, *C. aquatica, Stenotrophomonas sp*. and *B. pumilus*. Bacteria were grown overnight on Lysogeny broth (LB) plates at 30°C (*Stenotrophomonas sp*. and *B. pumilus*) and 37°C (*E. coli*, and *S. enterica*) from frozen glycerol stocks. The next morning a scoop of the bacterial lawn was inoculated in LB with antibiotics when required. Streptomycin 25 μg/mL (Calbiochem) was used for *E. coli* OP50 and grown for six hours on agitation at 200 rpm at 37°C (until an OD600 of 1.5-2.0). A volume of 100 mL of the bacterial cultures was seeded onto 60 mm NGM plates and allowed to dry overnight.

### NAD^+^ supplementation

1 mM NAD^+^ (Sigma) final concentration was added to liquid bacteria before seeding the NGM plates. *mec-4d* embryos obtained from synchronization of gravid adults were placed on the bacterial lawn 24 hours after they were seeded. Scoring of the neuronal integrity was done as explained below.

## Quantification of life history traits

### Developmental rate

60 mm NGM plates were seeded with 100 µL of desired bacteria previously cultured to an O.D.600 of 0.4. Thirty to forty 0-2 h post hatching L1 animals worms were placed per plate. Every 24 hours for three days the number of animals in each developmental stage was counted under a Nikon SMZ745T stereomicroscope with a Nikon G-AL 2x magnification.

### Pharyngeal pumping

30 adult animals (12 hours post L4) were observed under a stereomicroscope at high magnification. The number of pumping per minute of the animals was quantified using a manual counter.

### Defecation intervals

The time elapsed between defecation events in 30 adult animals (12 hours post L4) observed under a stereo microscope at high magnification was recorded using a chronometer.

### Locomotion pattern

10-15 adult nematodes were placed inside a bacterial lawn. Worm tracks were observed after one hour and therein followed for one more hour. If nematodes displayed movements with recurrent stops and turns, it was recorded as dwelling. If they showed rapid and incessant movement, it was recorded as roaming (13).

### Bacterial preference test

Food choice experiments of single and mixes of isolates were done in comparison with control bacteria *E. coli* OP50 and *E. coli* HT115. 60 mm NGM plates were prepared without antibiotics, and inoculated with bacteria at an absorbance of 0.4 nm on each side of the plate, generating a lawn of approximately 1 cm in diameter (30 µL). The plates were then left for 24 hours at room temperature to dry and the next day 30 worms in stage L4 previously fed with *E. coli* OP50 were placed in the border of the plate equidistant from each bacteria lawn. Animals were placed for 30 minutes on an antibiotic-free NGM plate to clean them of bacteria before placing them on the plates of preference. After being placed on the plate animals were quantified at 30 minutes and 1, 2, 3, 4, 12 and 24 hours.

### Occupancy in the bacterial lawn

60 mm NGM plates were seeded with 300 µL of a bacterial inoculum incubated at 37°C in agitation until O.D. 0.4 nm. Next day, 30 L4 animals previously fed on *E. coli* OP50 were placed on an antibiotic-free NGM plate to clean them of bacteria before being transferred by picking and placed in the center of the lawn. The number of worms inside and outside the lawn was quantified at 1, 2 and 3 hours.

### Neuronal integrity

Neurons with full-length axons were classified as AxW. Neurons with axons with a process connected to the nerve ring were classified as AxL, and those with axons that did not reach the bifurcation to the nerve ring were classified as AxT. The absence of the neuron, or neurons presenting only the soma, as well as neurons with axons conserving only the ventral projection were classified as Axϕ.

### Touch response

Animals were gently touched with an eyelash (17), in the anterior area over the second bulb of the pharynx. A positive response was recorded when animals moved backward as a result of being touched with the eyelash.

### Quantification of *acdh-1:: gfp* expression

Animals were mounted on 200 μm thick agarose pads with 1mM levamisole and visualized on an upright Eclipse Ti microscope (Nikon instruments) imaged with a 40x/0.75 N.A. objective and acquired with an EOS rebel t3i camera (Canon). Fluorescence was quantified using the software ImageJ, by cropping the region of interest (ROI) corresponding to individual animals, and taking the average intensity of pixels within each ROI.

### Statistical evaluation and experimental size

All experiments were done at least 3 times (three biological replicas, started in different days and from different starting plates). Each biological replica contained a triplicate (three technical replicas). Statistical evaluation was done by a one or two-way ANOVA with post-hoc analyses.

### Genomics

#### Extraction of bacterial DNA

Genomic DNA was extracted from cultures grown until an OD600nm between 0.4-0.6 nm using the Ultra Clean Microbial DNA Isolation Kit (MoBio Laboratories) according to the instructions of the manufacturer. Genomic DNA samples were quantified on a Nanoquant Tecan Infinite 200Pro.Genomic DNA samples from the bacterial isolates were treated with RNAse A (Qiagen) for 60 minutes at 37 °C to eliminate RNA remnants from the samples.

#### Genome sequencing and identification of bacteria

The genomic DNA samples of each bacterial isolate were sequenced at Genoma Mayor SpA employing the the TruSeq Nano DNA LT Kit (Illumina). A quality control of the sequenced reads of each sequenced sample was performed using the FastQC program. Next, readings were assembled using SPAdes. The quality of the genome assemblies was assessed with Quast (http://quast.bioinf.spbau.ru/). Genome assemblies were annotated using the RAST tool kit from PATRIC and deposited in BioProject database PRJNA876939. To identify the bacteria, genome sequences contigs were uploaded into the Type Strain Genome Server (TYGS, https://tygs.dsmz.de/) for taxonomy-based identification analysis. The resulting16S rRNA-based and Genome BLAST Distance Phylogeny (GBDP) -based phylogenetic trees and the digital DNA-DNA hybridization results table were retrieved. Form this analysis, the closest type strain to each isolate was determined. Next, the genomes sequences of the closest type strains were retrieved form GenBank and used to determine the average nucleotide identity (ANI) between the isolate’s genomes and their respective closest type strain using OrthoANIu (https://www.ezbiocloud.net/tools/ani).

## Metabolomics

### Generation of bacterial extracts

Bacteria were cultured in 2 mL of liquid LB at 30°C and 200 r.p.m overnight per ∼16 hours. The pre-incubated inoculum was placed in 30 mL of liquid LB in 50 mL Falcon tubes and grown at 30°C at 200 rpm until an OD600nm = 1. Bacterial cultures were centrifuged at 6.500 g for 5 minutes, an aliquot of 1 mL of the supernatant was stored at -20°C and the rest was discarded. The bacterial pellet was resuspended with 20 mL of a cold phosphate buffered saline solution (PBS: NaCl [137 mM], KCl [2.7 mM], Na_2_HPO_4_ [10 mM] and KH_2_PO_4_ [1.8 mM]). Pellets were centrifuged again and washed twice with 1 mL of cold PBS. After the washes the pellet was stored at -20°C. The pellet was resuspended with 600 µL of extraction buffer (acetonitrile and KH_2_PO_4_/NaH_2_PO_4_ 100 mM, pH 7.4) and mixed by vortexing for 30 seconds. Next, the tubes containing the resuspended pellet were placed in liquid nitrogen for 1 minute and then on ice until thawed (4°C). The vortex and frozen by liquid nitrogen steps were repeated two more times, and then the tubes were sonicated in an ultrasonic water bath (Bioruptor UCD-200, Diagenode) for 15 cycles of 30 seconds ON and 30 seconds OFF at maximum power. Then, the samples were centrifuged at 12.000 rpm for 10 minutes and the supernatants were recovered in new sterile tubes. The remaining pellet was extracted again as described and the supernatant resulting from the second extraction added to the first supernatant to optimize the recovery of metabolites. Supernatants were dried in a vacuum dryer (Savant) at 50°C for 2-3 hours depending on the number of samples. Once the samples were dried, they were stored at -20°C until metabolic profiling.

### Sample preparation for ^1^H NMR spectroscopy

The ^1^H NMR spectroscopy and multivariate data analysis were performed at the Platform for Structural and Metabolomic Biology (PLABEM), Rosario, Santa Fe, Argentina. Samples were randomized and reconstituted in 600 µL of 100 mM Na^+^/K^+^ buffer (pH 7.4) containing 0.005% sodium 3-trimethylsilyl-(2,2,3,3-2H4)-1-propionate (TSP) and 10% D2O. In order to remove any precipitate, samples were centrifuged for 10 minutes at 14,300g at 4°C. An aliquot of 500 µL of the centrifuged solution was transferred to a 5-mm NMR tube (Wilmad LabGlass).

### ^1^H NMR spectroscopic analysis of bacterial extracts

Water-suppressed ^1^H NMR spectroscopy was performed at 300 K on a Bruker 700 MHz spectrometer equipped with a 5-mm TXI probe (Bruker Biospin, Rheinstetten, Germany) using a standard 1D pulse sequence (noesygppr1d) (37, 38). The ^1^H NMR spectra were acquired using 4 dummy scans and 32 scans with 64 K time domain points and a spectral window of 20 ppm. The mixing time was set to 10 milliseconds, the data acquisition period to 2.228 seconds and the relaxation delay to 4 seconds. The free induction decays (FIDs) were multiplied by an exponential weighting function corresponding to a line broadening of 0.3 Hz.

### ^1^H NMR spectral processing

NMR spectra were processed in Matlab (version R2016b, The MathWorks). Spectra were referenced to TSP at 0.0 ppm, and baseline correction and phasing of the spectra were achieved using Imperial College written functions (provided by T. Ebbels and H. Keun, Imperial College London). Each spectrum was divided into integrated regions of equal width (0.04 ppm, bucket width). Non-informative spectral regions containing TSP and water signals were excluded. Each spectrum was then normalized by the probabilistic quotient method (39). Spectra alignment was made using the algorithm recursive segment-wise peak alignment (40), in user-defined windows and the full spectra matrix was exported.

### NMR Resonances integration

The resolution and integration of all resonances of the spectral dataset were performed using MCR-ALS as proposed by Pérez et al (41). Briefly, the spectral mode was manually segmented into 63 spectral subregions that were then analyzed throughout MCR-ALS. From the 63 integrated regions, 25 metabolites were assigned using public NMR databases (42), whereas 35 resolved curves could not be assigned. Unknown features were discarded and assigned features were selected to build the final matrix. In cases where more than one signal from a compound was integrated, the better resolved and less overlapped one was selected to build the matrix. The description of integrated regions and the number of considered components are detailed on **Table S1**. In cases where more than one signal per molecule was considered, integrated values were averaged.

### Multivariate data analysis

The full spectra matrix corresponding to single and mixed bacteria were imported to SIMCA 16 (Umetrics AB, Umeå, Sweden). Principal Component Analysis (PCA) was performed on the mean-centered and Pareto-scaled NMR dataset and score plots were analyzed. The absence of outliers was evaluated by Hotelling’s T2 ellipse at 95% confidence intervals.

To evaluate neuroprotection, an integrated peak matrix was imported to SIMCA. Data was mean centered and Unit Variance (UV) scaled. PCA was built for global data visualization and Orthogonal partial least squares discriminant analysis (OPLS-DA) was made to maximize the separation between bacterial groups as a function of neuroprotection (21). To ensure valid and reliable OPLS-DA models and to avoid over-fitting, 200 permutations were carried out Discriminant features between classes in OPLS-DA models were defined using a loading plot.

## One factor Statistical analysis

Peak integral matrix was imported to MetaboAnalyst 5.0 (MetaboAnalyst 5.0: narrowing the gap between raw spectra and functional insights). Volcano plots were used setting Fold Change = 2.0 and p < 0.05.

Accession number: The genomic data on the bacteria identified in this study has been deposited in BioProject database PRJNA876939.

## Figure Legends

**Figure S1.**
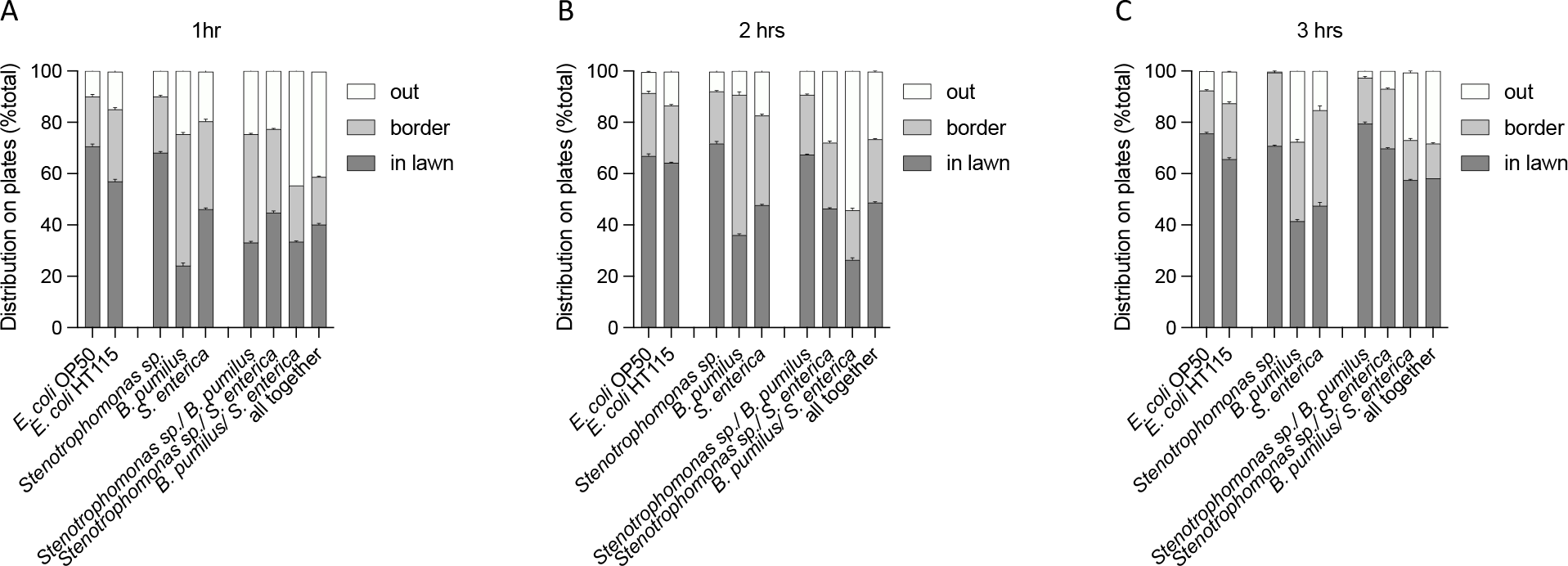
Percentage of animals inside, on the border and outside individual bacterial lawn or mixes after 1, 2 and 3 hours of placing them in the center of the plate.

**Figure S2.**
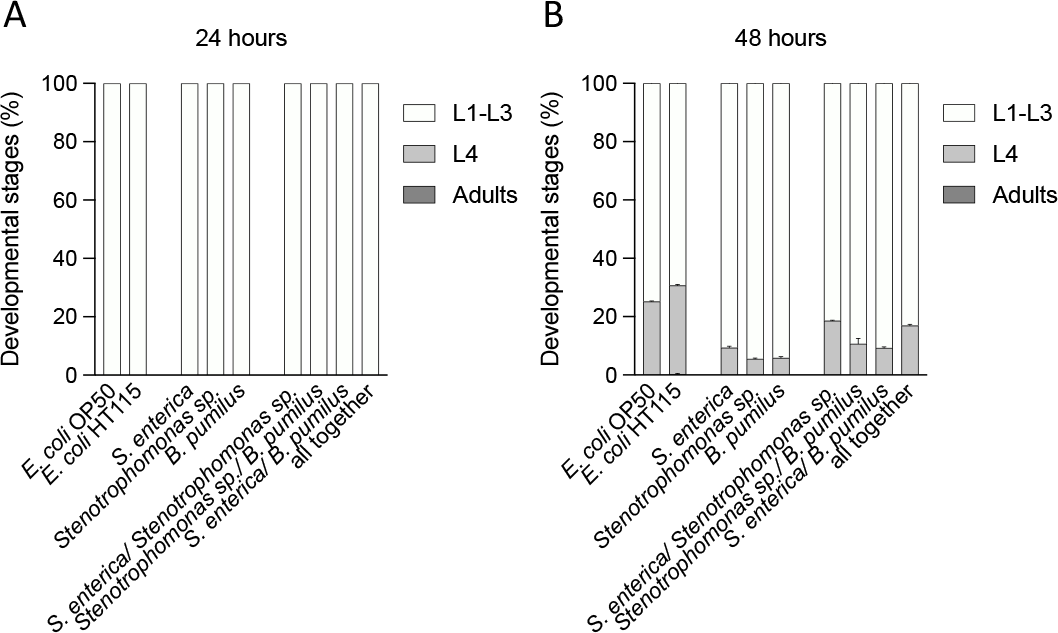
Percentage of animals in larval or adult stages or only adults at 24 (**A**) and 48 (**B**) hours after exposure to single bacteria or consortia.

**Figure S3.**
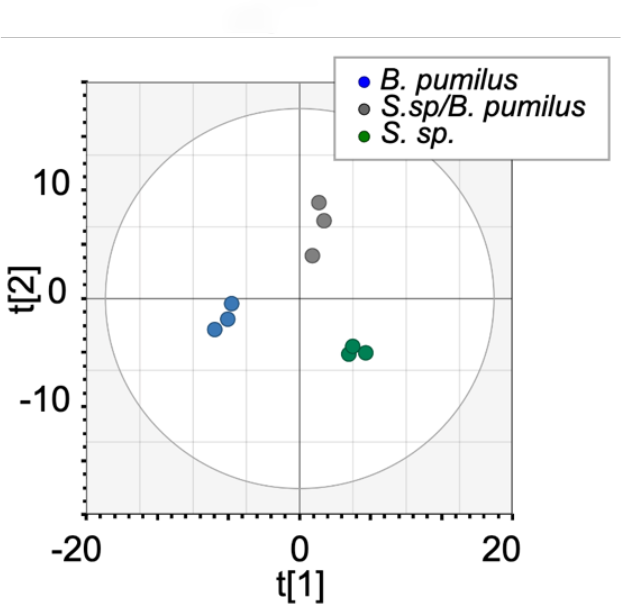
Principal Component (PC) scores plot derived from protective *B. pumilus* (blue) and nonprotective *Stenotrophomonas sp*. (green) and the mixture *B. pumilus/Stenotrophomonas sp*. (gray).

**Figure S4.**
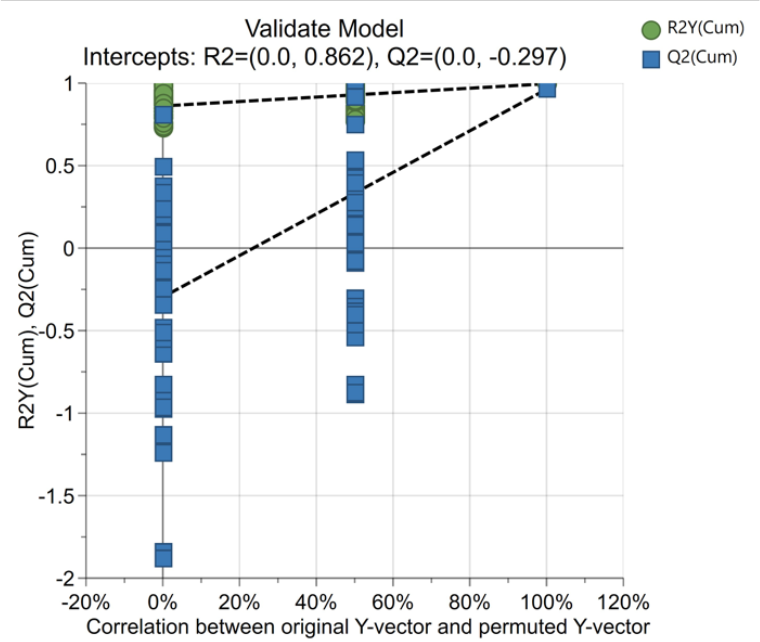
Validation of OPLS-DA model of protective *B. pumilus* and nonprotective *Stenotrophomonas sp*. and the mixture *B. pumilus/Stenotrophomonas sp*. by 200 permutations.

**Figure S5.**
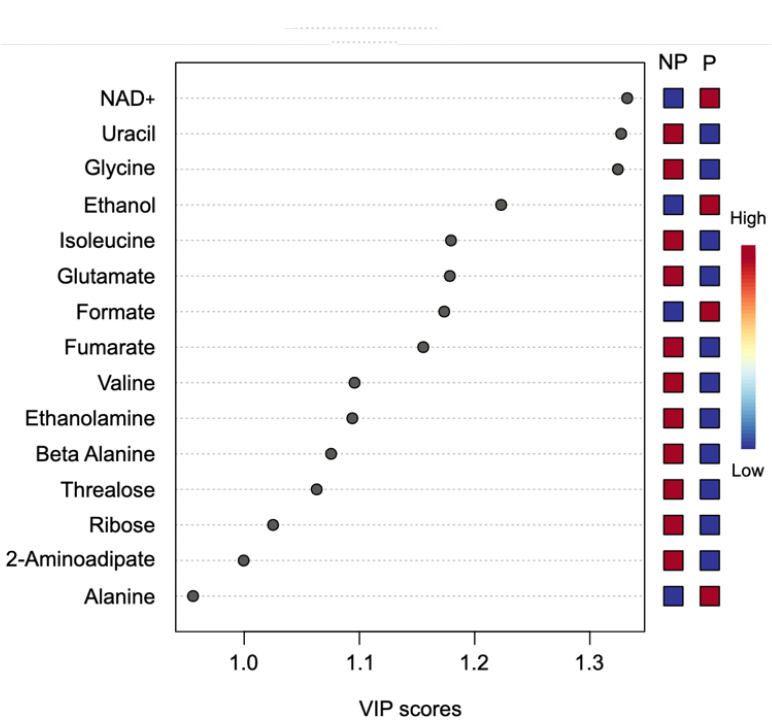
Variable importance in projection (VIP) plot displays the top 15 most important metabolite features identified by OPLS-DA. Colored boxes on right indicate relative concentration of corresponding metabolite for samples from protective (P) and nonprotective bacteria (NP).

**Table S1.**
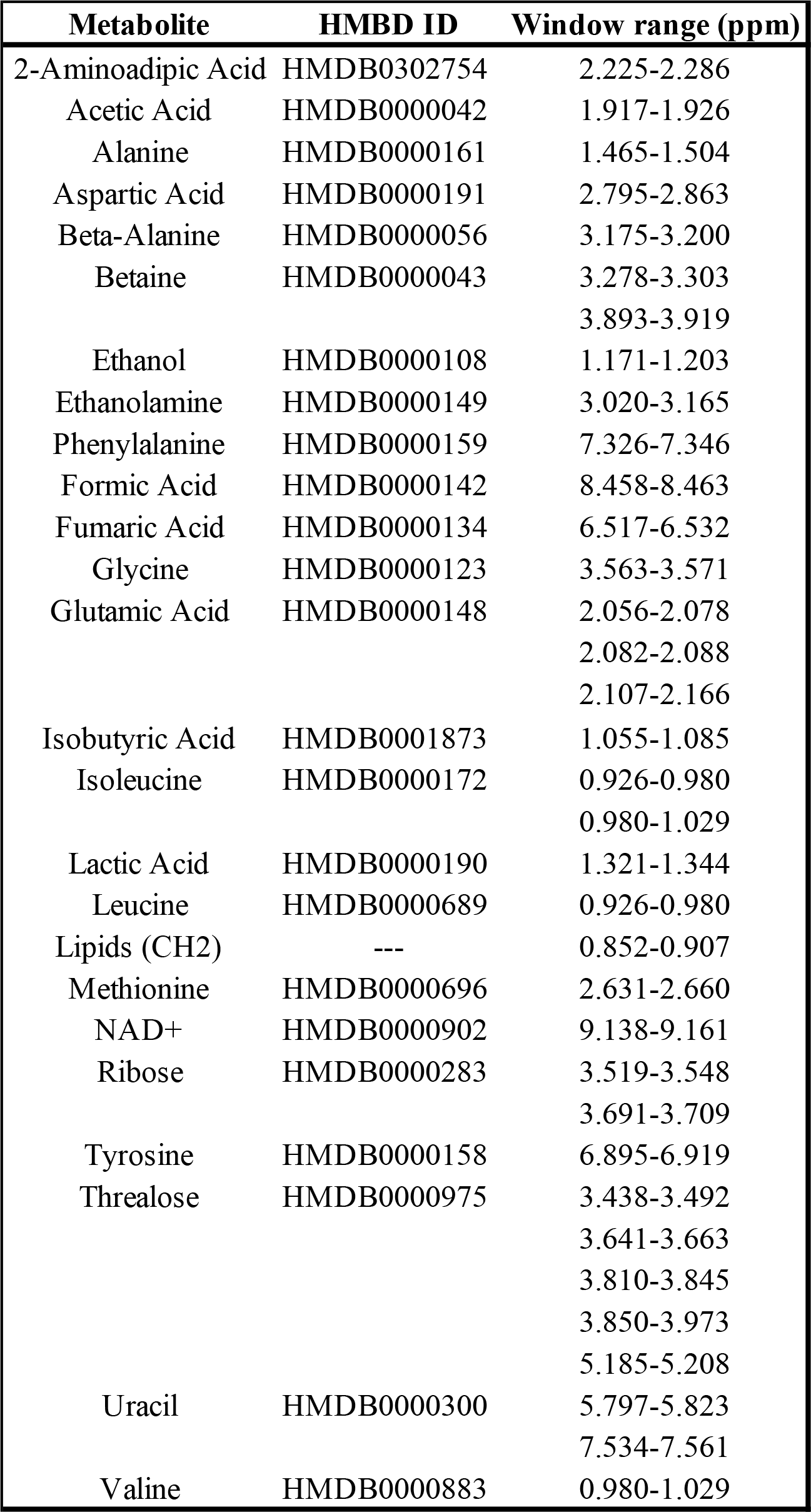

## Author contribution

Conceptualization SU, VG, AC

Methodology SU, PB, VG, MFP, JPC, AC

Investigation SU, PB, VG, MFP, LH, JPC, AC

Writing-Original Draft AC, MFP, VG, SU

Writing-Review and Editing SU, PB, VG, MFP, AC

Funding Acquisition AC

